# Alzheimer’s pathology is associated with dedifferentiation of functional memory networks in aging

**DOI:** 10.1101/2020.10.14.340075

**Authors:** Kaitlin E. Cassady, Jenna N. Adams, Xi Chen, Anne Maass, Theresa M. Harrison, Susan Landau, Suzanne Baker, William Jagust

**Author notes:** Corresponding author: Kaitlin Cassady, University of California Berkeley, Helen Wills Neuroscience Institute, 132 Barker Hall, Berkeley, CA 94720, USA.

## Abstract

In presymptomatic Alzheimer’s disease (AD), beta-amyloid plaques (Aβ) and tau tangles accumulate in distinct spatiotemporal patterns within the brain, tracking closely with episodic memory decline. Here, we tested whether age-related changes in the segregation of the brain’s functional episodic memory networks - anterior-temporal (AT) and posterior-medial (PM) networks - are associated with the accumulation of Aβ, tau and memory decline using fMRI and PET. We found that AT and PM networks were less segregated in older than younger adults and this reduced specialization was associated with more tau and Aβ in the same regions. The effect of network dedifferentiation on memory depended on the amount of Aβ and tau, with low segregation and pathology associated with better performance at baseline and low segregation and high pathology related to worse performance over time. This pattern suggests a compensation phase followed by a degenerative phase in the early, preclinical phase of AD.

## Introduction

The accumulation of beta amyloid (Aβ) plaques and neurofibrillary tau tangles is associated with episodic memory loss in both normal and pathological aging^1,2^, but the mechanisms underlying this association are not understood. Molecular and animal studies suggest that these pathologies spread through structurally and functionally connected brain regions^3–6^, with human neuroimaging studies indicating that patterns of tau deposition conform to large-scale brain networks in older adults (OA)^7–9^. Given that AD pathology starts to deposit in episodic memory networks, we investigated whether age-related changes in functional connectivity in these networks were associated with the accumulation of Aβ and tau.

Functional connectivity (FC) – the co-activation of brain regions within brain networks – reflects the brain’s large-scale network architecture. Brain networks specialized for different cognitive functions become dedifferentiated, or less segregated from each other, with older age^10^. In task-free functional magnetic resonance imaging (fMRI) studies, this progression is typically characterized by decreased within- and increased between-network FC at rest^11–15^. This decrease in network segregation leads to reduced specialization of neural networks and has been linked to OA’s worse performance relative to younger adults (YA) in several behavioral domains^11,13,16,17^. In contrast, other studies have found that less segregated networks are associated with *better* performance, suggesting that dedifferentiation may reflect greater plasticity or compensatory processes that occur during normal aging and neurodegeneration^18–24^. One potential reason for this discrepancy may be uncertainty about the molecular and pathological processes that drive network reconfiguration.

During the early stages of neurodegeneration, normal cognitive performance is often maintained despite neuronal loss, changes in network function, or the accumulation of neurodegenerative pathologies^25–27^. Such compensation is typically only evident in preclinical or mild cases of neurodegeneration, diminishing once the neurodegenerative pathology becomes too severe. There is also evidence that greater FC can be associated with either better or worse memory performance depending on disease severity. For instance, Van Hooren and colleagues found that greater FC between the default mode network and the dorsal attention network was associated with better memory in a cognitively normal group, but with worse memory in an MCI group^28^. One potential interpretation of these results is that although connectivity between different networks is beneficial early, it may fail to support compensation as pathology increases. In addition, it could provide a means for that pathology to spread.

Events encoded as episodic memories usually combine information about objects/items and scenes/context. Processing of these two types of information depends on distinct cortical pathways in the neocortex and medial temporal lobe (MTL) that converge in the hippocampus^29–31^. Object processing involves an anterior-temporal (AT) system that includes fusiform gyrus (FuG)/perirhinal cortex, inferior temporal gyrus (ITG), and amygdala. In contrast, scene processing relies on a posterior-medial system (PM) that includes retrosplenial cortex (RSC), precuneus, and parahippocampal cortex (PHC).

In vivo positron emission tomography (PET) studies have demonstrated that Aβ and tau accumulate in distinct regions within these two subnetworks in the aging brain^32^. Specifically, tau initially deposits in the transentorhinal region^33,34^ and appears to spread throughout the AT system in both healthy aging and AD, although it eventually affects the PM system as well. In contrast, Aβ deposition preferentially affects the PM system^32^. Previous work has shown that the AT and PM functional networks have distinct patterns of resting state FC with entorhinal cortex subregions in YA, and that such patterns predict the spatial topography and level of cortical tau deposition in cognitively normal OA^35,36^. However, it remains largely unknown whether these networks change with age and whether the accumulation of Aβ and tau is associated with changes in their modular organization and, consequently, memory decline.

There were two mains goals of this study: first, to investigate the effects of age, Aβ and tau on the intrinsic functional architecture of the AT and PM memory networks and second, to examine how relationships between pathology and network segregation affect episodic memory performance. To that end, we use resting state fMRI (rsfMRI) to measure the segregation of the AT and PM networks in cognitively healthy YA and OA. After examining the effect of age on network segregation, we then use PET measures of Aβ and tau deposition in OA to explore the relationship between these pathologies and segregation in the AT and PM networks. Finally, we assess the relationship between Aβ and tau, segregation, and episodic memory performance at baseline as well as change in performance over an average of 6 years in OA. We test three hypotheses: 1) AT and PM networks will be less segregated in OA compared to YA; 2) Given their distinct spatial topographies, increased tau in OA will be associated with less segregated AT networks whereas increased Aβ will be associated with less segregated PM networks, and 3) Network segregation in OA will interact with Aβ and tau pathology to predict episodic memory performance at baseline as well as change in performance over time.

## Results

### Study participants

Fifty-five YA (ages 18-35) and 97 cognitively normal OA (ages 60-93) were included in this study. All YA and OA participants underwent structural and resting state functional MRI. All OA additionally underwent tau-PET imaging with ^18^F-Flortaucipir (FTP), Aβ−PET with ^11^C-Pittsburgh compound-B (PiB), and standard neuropsychological assessments. All OA participants had a baseline MMSE score of ≥ 26 and no history of significant medial illnesses or medications that affect cognition. Demographic information for each age group is presented in Table 1.

**Table 1.** Cohort demographics

|  | YA (n = 55) | OA (n = 97) |
| --- | --- | --- |
| Age | 24.9 ± 4.4 (18-35) | 76.4 ± 6.1 (60-93) |
| Sex (M/F) | 28/27 | 36/61 |
| Education (Yrs) | 16.3 ± 2.0 | 16.8 ± 1.9 |
| APOE ε4 (C/NC) | N/A | 28/66 (3 N/A) |
| Global PiB DVR | N/A | 1.17 ± 0.24 (0.92-1.89) |
| AT FTP SUVR | N/A | 1.28 ± 0.20 (0.98-2.3) |
| PM FTP SUVR | N/A | 1.18 ± 0.12 (0.94-1.63) |
| Aβ +/- | N/A | 42/54 (1 N/A) |
| Tau +/- | N/A | 30/66 (1 N/A) |

### AT and PM networks are less segregated with older age

Using rsfMRI, we measured functional network segregation in the AT and PM networks in YA and OA. First-level ROI-to-ROI functional connectivity analysis was performed using the CONN toolbox^37^. For this analysis, we used 12 unilateral FreeSurfer ROIs that included amygdala, FuG and ITG as part of the AT network and RSC, PHC, and precuneus as part of the PM network (Figure 1). Semi-partial correlations were used for these first-level analyses to determine the unique variance explained by each ROI, controlling for the variance explained by all other ROIs entered into the same model. For each participant, the mean rsfMRI time series for each ROI was extracted/computed. Then, the cross-correlation of each ROI’s time course with every other ROI’s time course was computed, creating a 12 × 12 correlation matrix for each subject. Correlation coefficients were converted to z-values using Fisher’s r-to-z transformation^38^. The diagonal of the matrix was removed and negative correlations were set to zero^39^. Network segregation values were calculated as the difference in mean within-network FC and mean between-network FC divided by mean within-network FC:

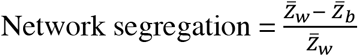

where 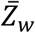 is the mean Fisher z-transformed correlation between ROIs within the same network and 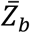 is the mean Fisher z-transformed correlation between ROIs of one network with all ROIs in the other network^13^. Thus, larger positive values for network segregation indicate that regions within a network (e.g. AT) have higher connectivity with each other compared to their connectivity with regions outside of the network (e.g. PM).

**Figure 1.**
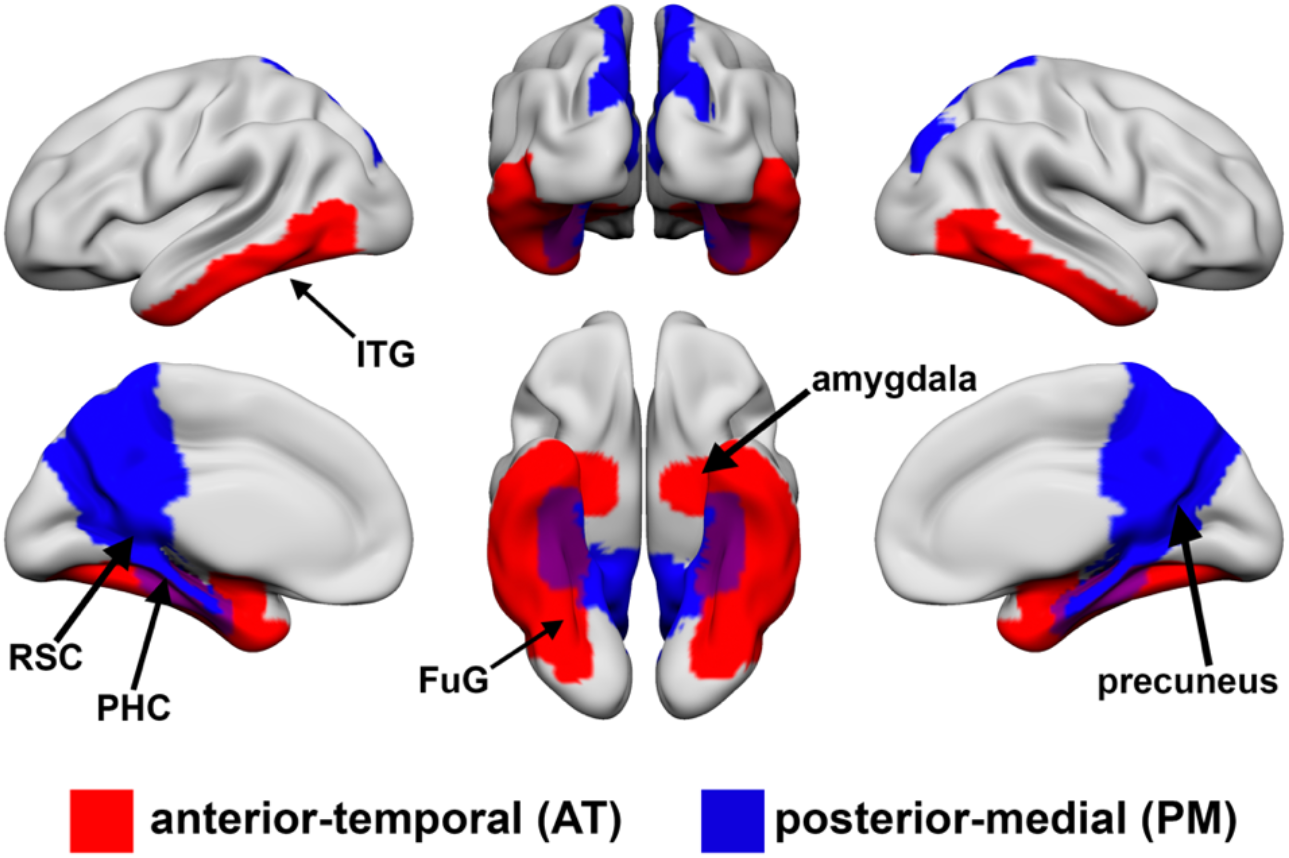
A priori defined regions of interest in anterior-temporal (AT; red) and posterior-medial (PM; blue) networks. AT regions include bilateral amygdala, fusiform gyrus (FuG)/perirhinal cortex, and inferior temporal gyrus (ITG). PM regions include bilateral retrosplenial cortex (RSC), parahippocampal cortex (PHC) and precuneus.

As hypothesized, OA exhibited decreased within-network (AT: Fig 2A, *t* = 6.7, *p* < 0.001; PM: Fig 2B, *t* = 3.1, *p* = 0.002) and increased between-network (AT: Fig 2C, *t* = 2.5, *p* = 0.01; PM: Fig 2D, *t* = 3, *p* = 0.003) functional connectivity and decreased segregation (AT: Fig2E, *t* = 4.9, *p* < 0.001; PM: Fig 2F, *t* = 4.3, *p* < 0.001) in the AT and PM networks compared to YA. Of particular importance, the relationship between age group and segregation was assessed across multiple analysis approaches related to matrix thresholding (i.e., inclusion of positive only vs. negative correlations), bivariate vs. semi-partial correlations, various network metrics of intersystem relationships (i.e., segregation, participation coefficient, and modularity), and network labeling (i.e., the regions included to define AT and PM networks). The age group differences in segregation were found to be robust in all instances (See Supplemental Figure S1).

**Figure 2.**
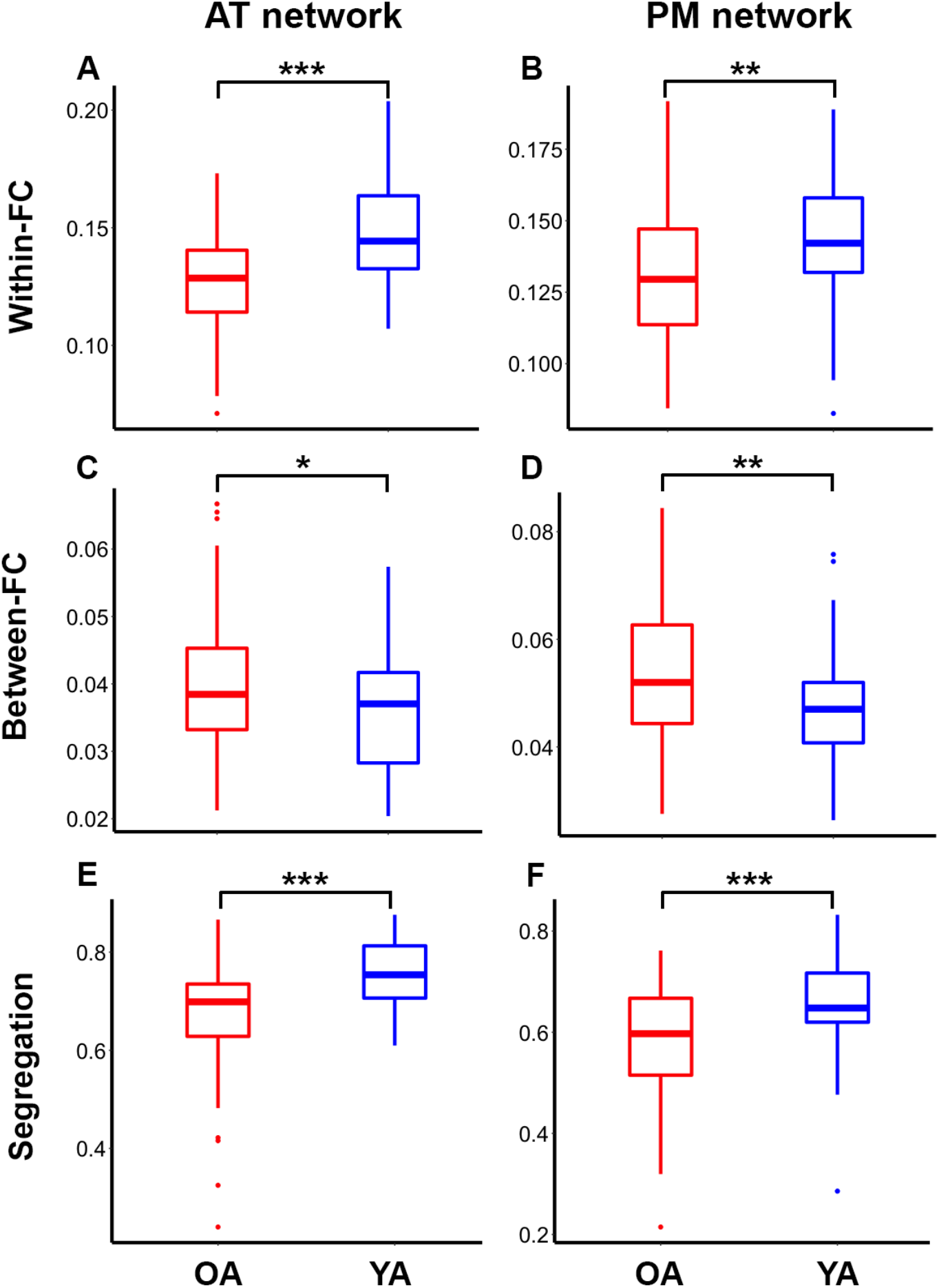
Anterior-temporal (AT; left) and posterior-medial (PM; right) networks are less segregated in older (red) relative to younger (blue) adults. OA show decreased within-network (**A** and **B**) and increased between-network (**C** and **D**) functional connectivity (FC), and decreased network segregation (**E** and **F**) compared to YA. \**p* < 0.05, \*\**p* < 0.01, and \*\*\**p* < 0.001.

### Tau relates to AT segregation and Aβ relates to PM segregation

Next, we examined the relationship between tau and Aβ pathology and network segregation in OA. Tau was quantified using the FTP tracer and data acquired at 80-100 minutes post-injection. The standardized uptake value ratio (SUVR) was calculated, and the weighted mean SUVRs for FreeSurfer-defined AT and PM ROIs (the same ROIs used for calculating segregation above) were calculated after partial volume correction (PVC)^40^. Aβ was quantified using the PiB tracer, and distribution volume ratio (DVR) was calculated. A global measure of PiB DVR was calculated across cortical FreeSurfer ROIs as described previously^41^, and a threshold of 1.065 was used to classify participants into Aβ− and Aβ+ groups^42^. *APOE* was not related to segregation (*ps* > 0.63); therefore, we did not control for *APOE* in these analyses.

To assess the relationship between tau and segregation, we first examined the relationship across all OA. We then split the group into Aβ− and Aβ+ subgroups to determine whether this relationship differed between OA with and without Aβ pathology. Increased FTP SUVR in AT regions was associated with decreased segregation in the AT network across all OA (Fig 3A, *r* = −0.25, *p* = 0.014). Splitting the group into Aβ− and Aβ+ subgroups, we observed the same relationship in the Aβ+ group (Fig 3B, *r* = −0.48, *p* = 0.002), but no significant relationship was observed in the Aβ− group (*r* = 0.10, *p* = 0.45). We did not find a significant relationship between global PiB DVR and AT network segregation across all OA (Fig 3C, *r* = −0.13, *p* = 0.23). Because the relationship between AT-tau and AT segregation appeared to be influenced by a few high-tau individuals, we performed a follow-up robust regression which is less affected by more extreme data points^43^. This analysis was performed using the ‘fitlm’ function with the ‘RobustOpts’ name-value pair in Matlab to create a model that limits the influence of outliers and heteroscedasticity. The relationship between tau and AT segregation was no longer significant across the whole group (*t* = 0.95, *p* = 0.34), but remained significant in the Aβ+ group (*t* = 3.1, *p* = 0.004).

**Figure 3.**
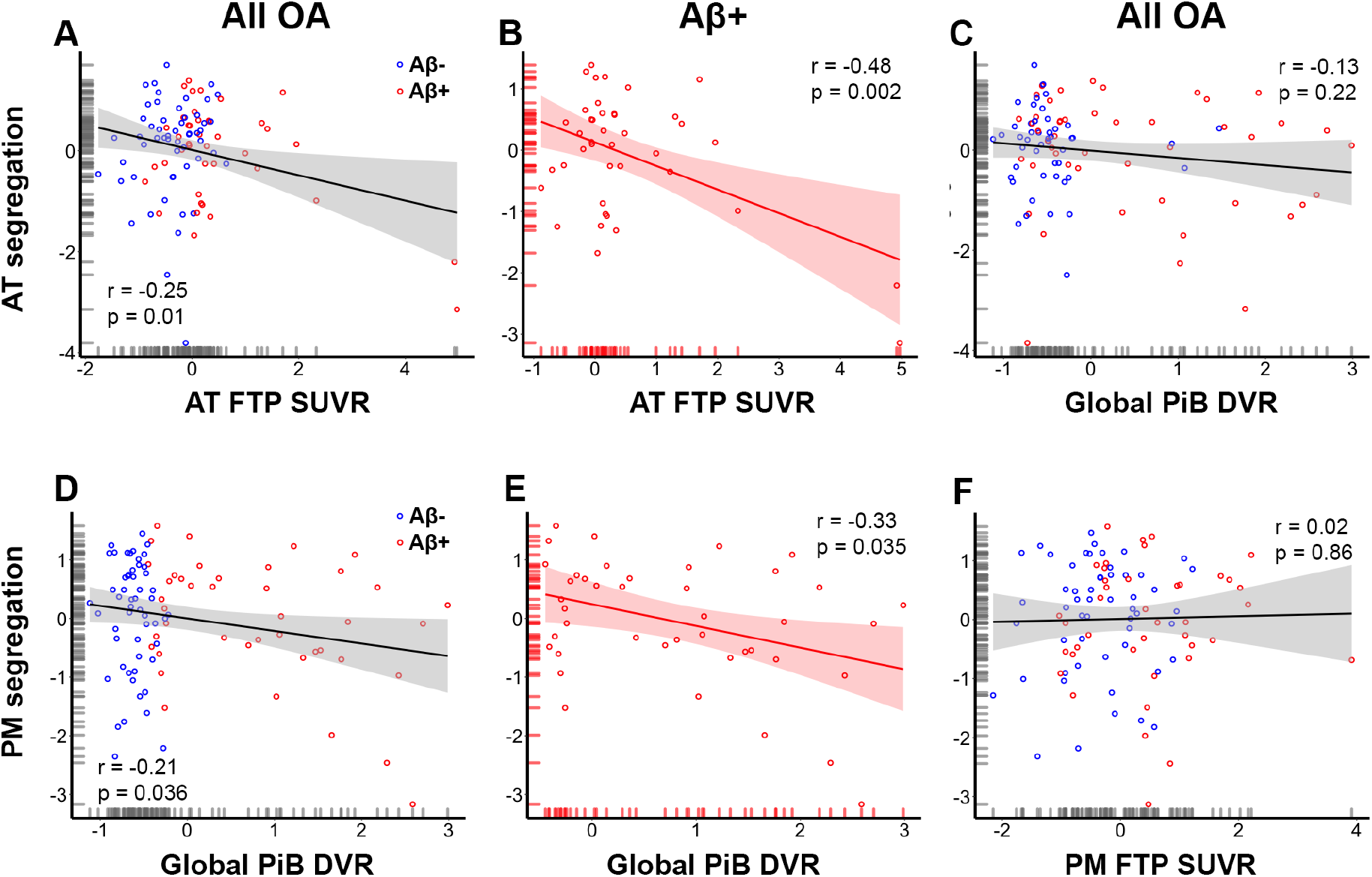
Tau and beta amyloid (Aβ) deposition are related to the segregation of anterior-temporal (AT; first row) and posterior-medial (PM; second row) networks, respectively. Less segregated AT networks are associated with increased tau in AT regions (**A**), particularly in Aβ+ OA (**B**), but are not associated with global Aβ (**C**). Less segregated PM networks are associated with increased global Aβ (**D**), particularly in Aβ+ OA (**E**), but are not associated with tau in PM regions (**F**). All regressions account for age and sex differences; The x and y axes reflect the residuals from the model.

We next examined the relationship between global PiB and network segregation. Increased global PiB DVR was associated with decreased segregation in the PM network (Fig 3D, *r* = −0.21, *p* = 0.036). Splitting the group into Aβ− and Aβ+ subgroups, we observed the same relationship in the Aβ+ group (Fig 3E, *r* = −0.33, *p* = 0.035) but no significant relationship in the Aβ− group (*r* = 0.014, *p* = 0.92). We did not observe a significant relationship between FTP SUVR in PM regions and PM network segregation (Fig 3F, *r* = 0.02, *p* = 0.86).

### AD pathology moderates the association between segregation and episodic memory

Cognitive performance was measured as a z-score of standardized neuropsychological tests in three different domains: episodic memory, executive function, and working memory. We then examined the relationship between network segregation and cognitive performance at baseline. Because our episodic memory composite measure included both object- and spatial-related memory domains, we computed a single segregation measure by averaging the AT and PM segregation values (follow-up analyses showed essentially the same results for AT and PM networks; See Supplemental Figure S2).

We performed a multiple regression to assess the effects of segregation and Aβ−status on episodic memory performance in OA, controlling for age, sex, and education. We observed main effects of segregation (*t* = 2.9, *p* = 0.005), Aβ−status (*t* = 2.5, *p* = 0.013), and age (*t* = 2.7, *p* = 0.008) on episodic memory. This analysis also revealed a significant interaction between Aβ−status and segregation on episodic memory performance (*t* = 2.5, *p* = 0.014). Specifically, less segregated networks were associated with better performance in Aβ− OA (Fig 4A, *r* = −0.40, *p* = 0.004) whereas segregation was not associated with performance in Aβ+ OA (Fig 4B, *r* = 0.14, *p* = 0.40). Table 2 reports the results of this regression.

**Figure 4.**
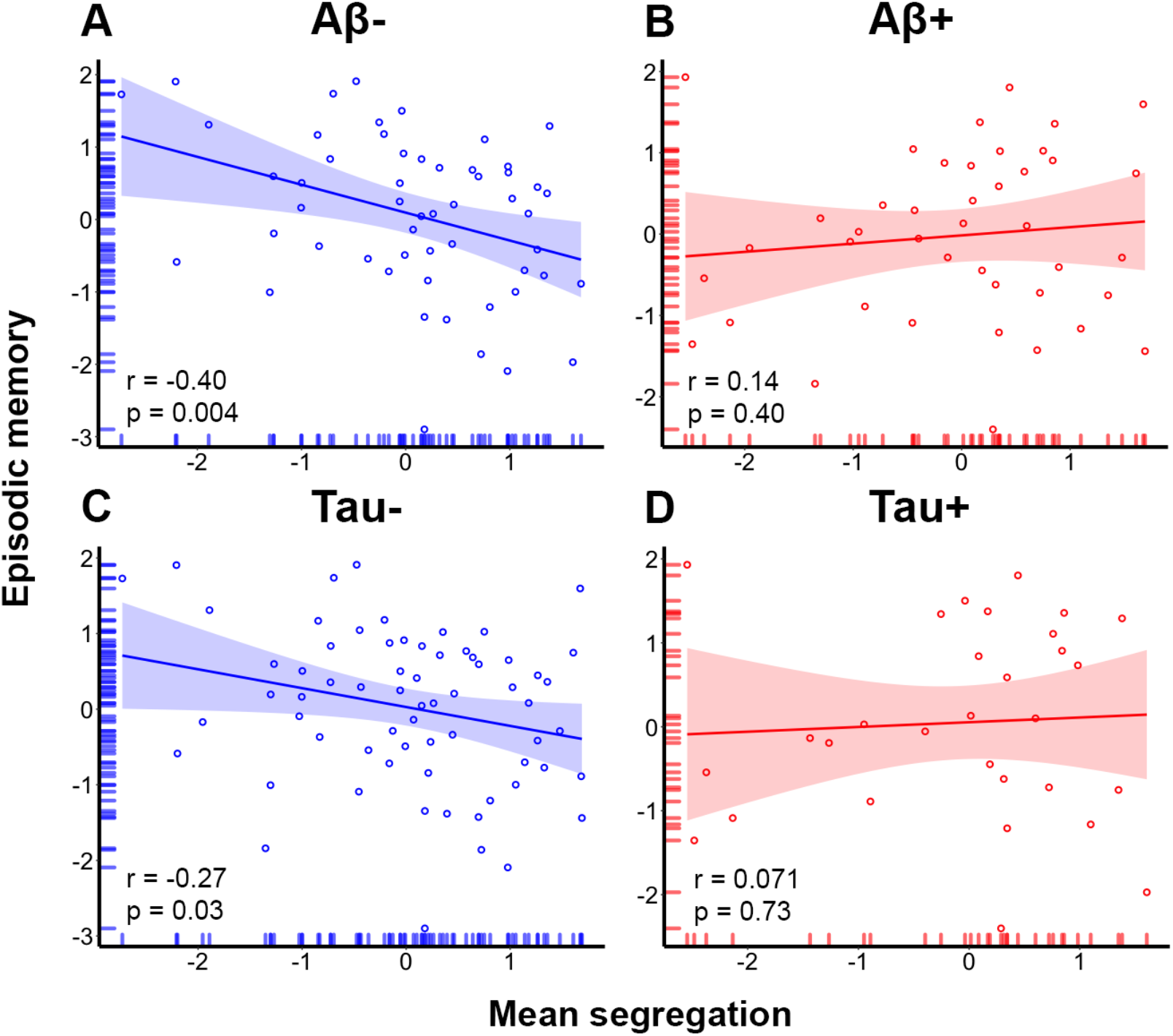
Alzheimer’s disease pathology moderates the association between mean network segregation and episodic memory performance. (**A**) Less segregated networks are associated with better performance in Aβ− OA whereas **(B)** segregation is not associated with performance in Aβ+ OA. (**C**) Similarly, less segregated networks are associated with better performance in Tau-OA whereas **(D)** segregation is not associated with performance in Tau+ OA.

**Table 2.** Multiple regression results for mean segregation and its interaction with Aβ−status predicting episodic memory at baseline.

| Predictor | t | p |
| --- | --- | --- |
| Age | -2.7 | 0.008 |
| Aβ-status | -2.5 | 0.013 |
| Sex | 0.86 | 0.39 |
| Education | 1.3 | 0.2 |
| Segregation | -2.9 | 0.005 |
| Aβ-status x Seg | 2.5 | 0.014 |

To additionally explore the effects of segregation and tau-status on episodic memory performance, we calculated “tau positivity” as the mean SUVR in a Braak_III-IV_ composite ROI (cut-off of 1.26) that included regions from both AT (amygdala, FuG, and ITG) and PM (PHC and RSC) systems^32,44^ as well as other regions that accumulate tau in the progression from aging to AD. We again found main effects of segregation (*t* = 2.1 *p* = 0.043) and age (*t* = 2.5, *p* = 0.014), but not tau-status (*t* = 1.4, *p* = 0.16) on episodic memory. Although there was not a significant interaction between tau-status and mean segregation on performance, (*t* = 1.5, *p* = 0.14) we did find that less segregated networks were associated with better performance in tau-OA (Fig 4C, *r* = −0.27, *p* = 0.03) whereas segregation was not associated with performance in tau+ OA (Fig 4D, *r* = 0.071, *p* = 0.73). Table 3 reports the results of this follow-up regression.

**Table 3.** Multiple regression results for mean segregation and its interaction with Tau-status predicting episodic memory at baseline.

| Predictor | t | p |
| --- | --- | --- |
| Age | -2.5 | 0.014 |
| Tau-status | -1.4 | 0.16 |
| Sex | 0.75 | 0.45 |
| Education | 1.2 | 0.22 |
| Segregation | -2.9 | 0.043 |
| Tau-status x Seg | 2.5 | 0.14 |

As control analyses, we also examined the relationships between segregation, Aβ−status, and baseline working memory and executive function. We again observed a significant interaction between Aβ−status and mean segregation on executive function (*t* = 2.21, *p* = 0.03) such that less segregated networks were associated with better executive function in the Aβ− OA (*r* = −0.35, *p* = 0.011) whereas there was no significant association in the Aβ+ OA (*r* = 0.08, *p* = 0.63). There were no significant relationships between segregation and working memory performance, nor was there an interaction between Aβ−status and segregation on performance (*ps*>.41).

Including tau-status in place of Aβ−status, we did not observe a significant interaction between tau-status and mean segregation on executive function, *t* = 1.5, *p* = 0.13. However, we did find that less segregated networks were associated with better executive function in the tau-OA (*r* = −0.28, *p* = 0.03) whereas there was no significant association in the tau+ OA (*r* = 0.08, *p* = 0.71). There were no significant relationships between segregation and working memory performance in either group, nor was there an interaction between tau-status and segregation on performance (*ps*>.67).

### Baseline segregation predicts longitudinal memory decline

To examine the effect of segregation and Aβ and tau on change in memory performance over time, our longitudinal analyses included participants that had at least two neuropsychological testing sessions (one of which was near the time of the resting state scan). Importantly, all participants had resting state fMRI, PET, and neuropsychological data at the same time point (i.e., “baseline”), which is the time point we used to assess all cross-sectional relationships. Eighty-six of 97 OA participants had longitudinal cognitive data (≥ 2 testing sessions. These participants had between 2 and 13 testing sessions (mean, 6.1 ± 3.1) over a period of 1 to 13 years (mean, 6.1 ± 3.5) with an average interval of 1.3 ± 0.6 years between sessions. Sixteen participants had only retrospective data. Figure 5A displays each participant’s trajectory in longitudinal episodic memory performance over time.

**Figure 5.**
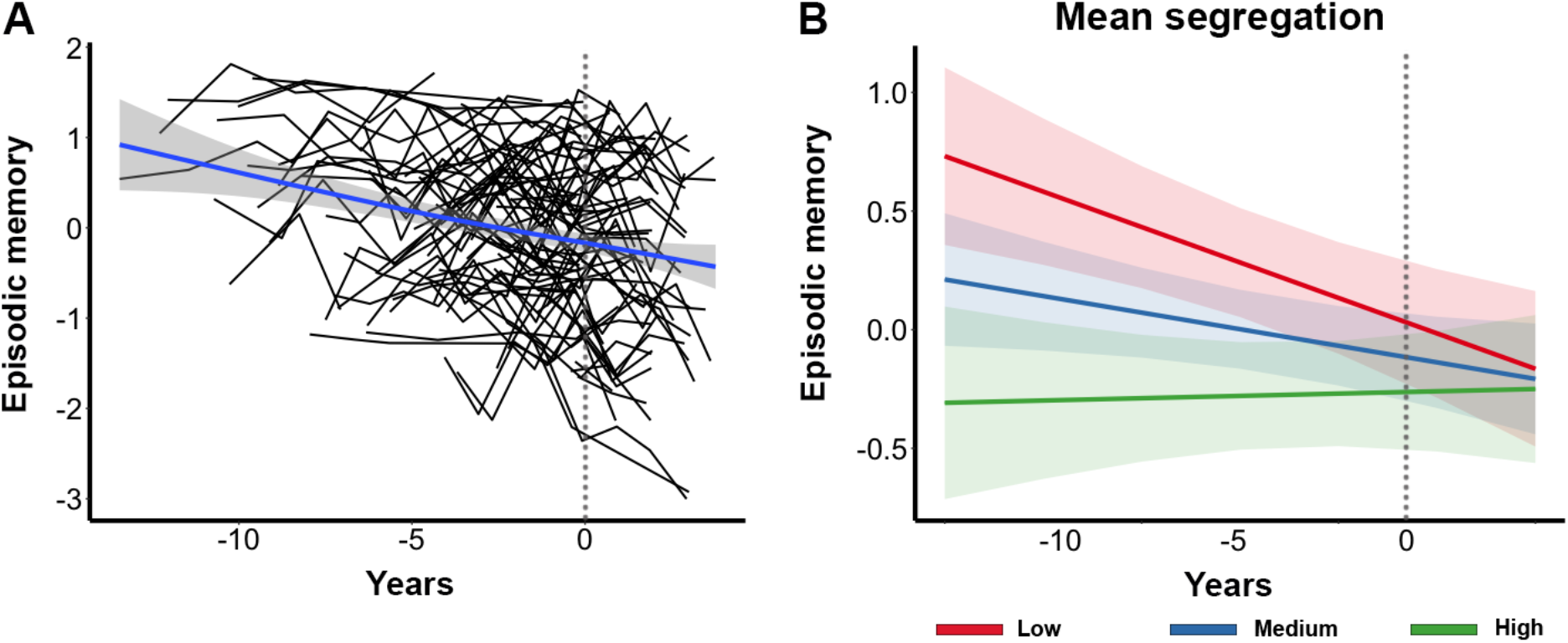
Relationship between network segregation and longitudinal episodic memory change. (**A**) Individual participant trajectories in longitudinal episodic memory change over time. Each black line represents one participant. The blue trendline reflects the participant average change in memory over time. The dotted gray line (at X = 0) represents the “baseline” time point in each plot. (**B**) Plot of estimated curves for three groups with different baseline network segregation (low, medium, and high) and episodic memory outcomes over time. Note that segregation was modeled as a continuous variable but is shown as a categorical variable for illustration purposes only. Lower baseline segregation was associated with a steeper decline rate in episodic memory over time.

Longitudinal cognitive measures were modelled using linear mixed-effects regression with a random intercept and slope using the lme4 package in R v3.6.3 (www.r-project.org). In order to examine the relationship between baseline segregation, baseline Aβ and tau, and change in cognitive performance in one model, the model included two-way interactions between baseline segregation and time, global Aβ and time, and tau and time. We report the results using continuous measures of global Aβ and Braak_III-IV_ tau as they retain more statistical power in the model. The results were very similar whether we used Braak_III-IV_ tau, AT-tau (Table S1) or PM tau (Table S2) and whether we used dichotomous (Table S3) or continuous Aβ and tau in the model. All models were adjusted for age, sex, and education. Segregation was a continuous variable in the model but is displayed graphically using tertiles.

We found that individuals with lower segregation at baseline showed a steeper decline rate in episodic memory over time (β = 0.08, SE = 0.03, *p* = 0.02; Figure 5B; Table 3). We also found that more tau at baseline was associated with a steeper decline rate in memory over time (β = −0.15, SE = 0.04, *p* = 0.002). There was no interaction of Aβ and time predicting memory change (β = −0.003, SE = 0.04, *p* = 0.94). To examine whether Aβ or tau moderated the effect of segregation on cognitive change, follow-up analyses included the same factors in addition to three-way interactions between baseline segregation, baseline Aβ and tau, and time. These analyses did not show a significant three-way interaction between segregation, tau and time (β = 0.03, SE = 0.05, *p* = 0.58) nor between segregation, Aβ, and time (β = −0.04, SE = 0.04, *p* = 0.34) on memory change. As a control analysis, we also examined change in working memory and executive function performance over time using the same model (not including three-way interactions). Baseline segregation was neither associated with longitudinal change in working memory (β = - 0.02, SE = 0.04, *p* = 0.57) nor executive function (β = 0.02, SE = 0.03, *p* = 0.66).

**Table 3.**
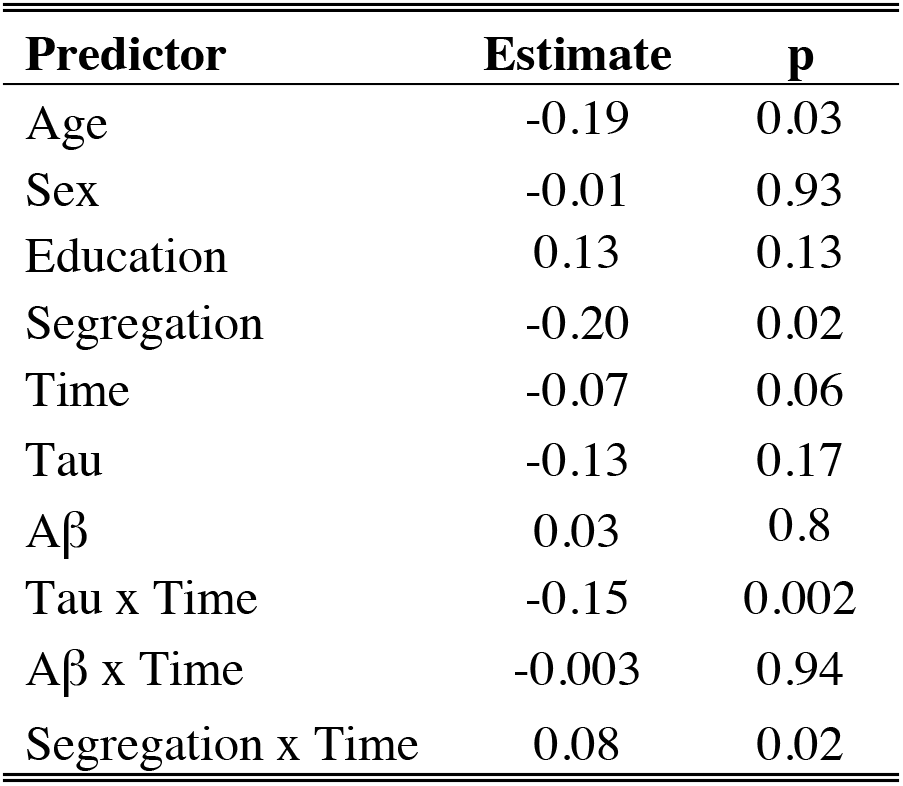
Linear mixed model results for segregation and pathology predicting longitudinal episodic memory change.

## Discussion

The goal of this study was to investigate the effects of Aβ and tau on the intrinsic functional architecture of episodic memory networks and episodic memory ability in cognitively normal OA. OA showed reduced segregation of AT and PM networks compared to YA. This effect was driven by reduced within-network FC and increased between-network FC between the two systems. Increased tau in AT regions was associated with a less segregated AT network, whereas increased global Aβ was associated with a less segregated PM network, demonstrating a regional dissociation of these AD pathologies to the large-scale organization of the AT and PM systems. Finally, less segregated networks were associated with better memory ability at baseline in OA with low levels of AD pathology but with a steeper decline in memory performance over time, independent of baseline pathology. These findings suggest there may be different phases in the association of brain network integrity and memory ability that depend on the degree of AD pathology.

We interpreted our findings based on a model that includes both age- and AD pathology-related effects. We found that age was associated with changes in the functional segregation of the AT and PM resting state networks. This finding is consistent with studies of age-related neural dedifferentiation demonstrating that older age is associated with less distinct neural activation patterns^10,11,45–48^ and, more recently, with less distinct large-scale resting state networks^11–15,49,50^. While a majority of these prior studies explored the organization of the brain’s canonical resting state networks (e.g., the default mode, fronto-parietal and cingulo-opercular networks), we demonstrated a robust age effect in two neural networks that are associated with episodic memory and AD pathology. Recent work from our laboratory also showed that less differentiated activation in AT and PM regions during an object/scene discrimination task was associated with more tau deposition^32^. These findings, in conjunction with the present results, suggest a neuropathological correlate of dedifferentiation in the episodic memory system.

We found that the modular organization of the AT and PM brain networks was selectively vulnerable to tau and Aβ deposition. Specifically, increased tau in AT regions was associated with less segregated AT networks, but was not associated with PM network segregation. In contrast, increased cortical Aβ was associated with less segregated PM networks, but was not associated with AT network segregation. Since between-network FC was the same in the AT and PM networks, these results indicate that this relationship was driven by within-network FC. This is consistent with previous investigations that have demonstrated relationships between within-network FC and AD pathology^8,35,51^. The findings of a double dissociation between AD pathology and network segregation are in accordance with previous work from our lab demonstrating differential selective vulnerability to these two networks participating in episodic memory function. Specifically, Maass et al. (2019) showed that tau deposits mainly in AT regions, resulting in object discrimination deficits, whereas Aβ deposits preferentially in PM regions, resulting in impaired scene discrimination^32^.

There is no agreement on a precise region where Aβ deposition begins, and existing data suggest that this pathology appears multifocally and quickly accumulates throughout most of association cortex^52–55^. For example, while we used a global measure of cortical Aβ to define positivity, Aβ in the PM network is highly correlated with this measure as are most regions throughout the brain^56^. In contrast, tau initially deposits in the entorhinal cortex and then progresses in a distinct spatiotemporal pattern first to anterior temporal and limbic regions and then throughout association cortex^52,57,58^. Cellular and molecular data reveal that tau can spread trans-synaptically and in relation to neural activity^5,6,59,60^, suggesting that this specific AD pathology accumulates through the brain along neural connections. The idea that large-scale brain network connectivity may underlie the spatiotemporal patterns of AD pathology has support from other laboratories. For instance, Franzmeier et al. found that canonical network regions with higher FC showed higher covariance of tau deposition^7^. In addition, Jacobs et al. found that Aβ facilitated the spread of tau from the hippocampus to the posterior cingulate via structural connectivity^61^, and Adams et al. reported that FC of the entorhinal cortex was related to Aβ−facilitated neocortical tau deposition^31^. Previous data suggesting preferential involvement of the AT network by tau^32^, along with these results showing dedifferentiation, raise the possibility that tau may spread from the AT to the PM network as these networks become less segregated.

Our results revealed complex interactions between segregation, Aβ and tau pathology, and memory performance. Our cross-sectional data showed that AD pathology moderated the relationship between segregation and baseline performance. Specifically, less segregated networks were associated with better performance in OA with low levels of pathology but not in those with high levels of pathology. Additionally, our longitudinal results revealed that less segregated networks and more tau at baseline independently predicted a steeper decline in memory performance over time. These findings are consistent with previous studies demonstrating that neurodegenerative pathology interacts with FC to influence performance^28,62^. For instance, Lin et al. (2020) showed an interactive effect of Aβ deposition and FC on cognition such that increased FC between left middle frontal gyrus and a memory encoding network was associated with better attention/processing speed and executive function in those with low levels of Aβ but with worse function in those with high levels of Aβ^62^.

Overall, our findings may suggest different phases in the long-term interaction of network segregation and AD pathology on episodic memory ability. OA with low pathology may compensate, either for normal aging processes or for the start of AD pathology, by increasing communication between AT and PM networks. As functionality in one system declines, recruiting the other system may help performance. Over time, however, this increased between-network FC in the context of increasing pathology could become detrimental, as well as providing a route for AD disease pathology to spread from one network to the other, leading to more decline in memory ability. Based on this model, it is likely that AT and PM networks continue to de-differentiate over time, especially in the transition phase from cognitively normal to cognitively impaired. This would further the spread of AD pathology, eventually resulting in the hallmark episodic memory impairments observed in MCI and AD. Future studies that include patient data as well as longitudinal measures of FC, AD pathology, and memory function are crucial in testing this hypothesis.

The cross-sectional PET and MRI data limit our interpretations of causality as well as long-term changes in this study. Although it is possible that Aβ and tau spread lead to disruptions in large-scale network FC (rather than the reverse), several studies suggest that Aβ and tau propagation is a multifactorial process that depends on both neural connectivity and regional vulnerability^6,8,60^. Hence, the relationship between Aβ and tau and FC is likely bidirectional such that age-related disruptions in network FC guide pathology spread and this, in turn, leads to further changes in the network architecture. Longitudinal designs are critical in determining the order of age-related changes as well as elucidating the sequence of neural events leading to episodic memory decline. Another limitation of this study was that the longitudinal cognitive data included different numbers of timepoints before and after the “baseline” timepoint for different participants. This design feature complicates our interpretation of the longitudinal effects of segregation and Aβ and tau on performance because our analyses were both retrospective and prospective. However, this design feature allowed us to examine memory change over a longer period of time (average of ~6 years) compared to many previous studies^63–65^. Furthermore, we were able to include more participants from our sample with longitudinal data using this design. Longitudinal studies are often unable to observe any significant change in cognition in OA given the relatively short time periods of observation^66,67^. We believe that having a greater number of timepoints for more participants outweighs the disadvantage of this design feature.

Taken together, our data support a model whereby network dedifferentiation performs a neural compensatory function which fails over time as AD pathology accumulates. The effect of network dedifferentiation on episodic memory ability is helpful to performance when pathology levels are low but is harmful to performance over time as pathology presumably spreads. This research provides an important step in elucidating the neural mechanisms associated with episodic memory decline in healthy and pathological aging. By studying this episodic memory system in healthy OA, we can advance our understanding of healthy aging and its similarities to and differences from pathological aging, which could serve as a crucial building block for the early detection of AD.

## Methods

### Participants

Fifty-five YA (age 18-35) and 97 cognitively normal OA (age 60+) enrolled in the Berkeley Aging Cohort Study (BACS) were included in this study. All YA and OA participants underwent structural and resting state functional MRI. All OA additionally underwent tau-PET imaging with ^18^F-Flortaucipir (FTP), Aβ−PET with C-Pittsburgh ^11^Compound-B (PiB), and a standard neuropsychological assessment. Eligibility requirements included that all participants had a baseline MMSE score of ≥ 26. We also excluded any participants with a history of significant neurological disease (e.g., stroke, seizure, loss of consciousness ≥ 10 minutes), or any medical illness that could affect cognition, history of substance abuse, depression, or contraindications to MRI or PET. All study procedures were reviewed and approved by the Institutional Review Boards of the University of California Berkeley and Lawrence Berkeley National Laboratory (LBNL). All participants provided written informed consent for their involvement in this study.

### Neuropsychological assessment

All OA participants in the BACS undergo neuropsychological testing to measure cognitive performance related to verbal and visual memory, working memory, processing speed, executive function, language, and attention. In this study, composite scores were calculated to measure three specific cognitive domains: episodic memory, working memory, and executive functioning. The tests for episodic memory included the California Verbal Learning Test (CVLT) immediate and long delay free recall totals as well as the Visual Reproduction (VR) immediate and delay recall totals. Working memory was assessed with the Digit Span total score. Tests for executive function included Trail Making test B minus A, Stroop number correct in 1 min, and Digit Symbol total. For episodic memory and executive function, the composite scores were produced by calculating the average z-score of the tests included in each domain. Please refer to Harrison et al. (2019) for more details of the procedure^44^.

To examine change in memory performance over time, our longitudinal analyses included participants (from the 97 OA sample) that had (1) at least one resting state fMRI scan, (2) at least one Aβ and one tau PET scan (both near the time of the resting state scan), and (3) at least two neuropsychological testing sessions (one of which was near the time of the resting state scan). Critically, all participants had resting state fMRI, PET, and neuropsychological data at the same time point, which is the time point we used to assess all cross-sectional relationships (i.e., “baseline” time point). The additional neuropsychological session(s) could be either before or after the baseline time point (or both), depending on the participant. Eighty-six of 97 OA participants had longitudinal cognitive data (≥ 2 testing sessions. These participants had between 2 and 13 testing sessions (mean, 6.1 ± 3.1) over a period of 1 to 13 years (mean, 6.1 ± 3.5) with an average delay of 1.3 ± 0.6 years between sessions.

### MRI data acquisition

All YA and OA participants underwent structural and functional MRI acquired on a 3T TIM/Trio scanner (Siemens Medical System, software version B17A) using a 32-channel head coil. First, a whole-brain high resolution T1-weighted volumetric magnetization prepared rapid gradient echo image (MPRAGE) structural MRI scan was acquired with the following parameters: voxel size = 1 mm isotropic, TR = 2300 ms, TE = 2.98 ms, matrix = 256 × 240 × 160, FOV = 256 × 240 × 160 mm^3^, sagittal plane, 160 slices, 5 minute acquisition time. This was followed by a rsfMRI scan that was acquired using T2*-weighted echo planar imaging (EPI) with the following parameters: voxel size = 2.6 mm isotropic, TR = 1067 ms, TE = 31.2 ms, FA = 45, matrix = 80 × 80, FOV = 210 mm, sagittal plane, 300 volumes, anterior to posterior phase encoding, ascending acquisition, 5 minute acquisition time. A multiband acceleration factor of 4 was used to acquire whole-brain coverage at high spatial resolution by acquiring 4 slices at the same time^68,69^. During the rsfMRI scan, participants were instructed to remain awake with their eyes open and focused on the screen, which displayed a white asterisk on a black background.

As part of the standard PET processing pipeline, a whole-brain high resolution T1-weighted volumetric MPRAGE scan was acquired for each participant on a Siemens Magnetom Avanto scanner at LBNL with the following parameters: voxel size = 1 mm isotropic, TR = 2110 ms, TE = 3.58 ms, flip angle = 15°, sagittal slice orientation. These data were used for PET coregistration and to parcellate the brain for PET data analysis.

### PET data acquisition

All OA participants underwent PET scanning at LBNL using a Biograph PET/CT Truepoint 6 scanner (Siemens, Inc.) with CT scans performed for attenuation correction prior to each emission acquisition and radiotracers synthesized at the LBNL Biomedical Isotope Facility. Tau deposition was measured using ^18^F-Flortacipir (FTP) with data binned into 4 × 5 minute frames from 80-100 minute post-injection^35,44,70^. Aβ was measured using ^11^C-Pittsburgh Compound B (PiB), with data acquired across 35 dynamic frames for 90 minutes post-injection (4 × 15, 8 × 30, 9 × 60, 2 × 180, 10 × 300, and 2 × 600 seconds). All PET images were reconstructed using an ordered subset expectation maximization algorithm, with attenuation correction, scatter correction, and smoothing using a Gaussian kernel of 4 mm.

### MRI processing

Structural scans (3T) were processed with FreeSurfer to derive regions of interest (ROIs) in each subject’s native space using the Desikan-Killany atlas. The structural images were also segmented into gray matter (GM), white matter (WM), and cerebrospinal fluid (CSF) using Statistical Parametric Mapping software (SPM12; Wellcome Trust Centre for Neuroimaging, London, UK) (default parameters). RsfMRI data were preprocessed using SPM12 and FreeSurfer (v5.3.0). Preprocessing included slice time correction, realignment, coregistration to the T1 image, and outlier volume detection. All functional images were first corrected for differences in slice time acquisition using SPM12. Functional images were then realigned to the first volume, and coregistered to the T1 image. Outliers in average intensity and/or scan-to-scan motion were identified using the artifact detection toolbox (ART; http://www.nitrc.org/projects/artifact_detect) using a conservative movement threshold of >0.5 mm/TR and a global intensity z-score of 3. Outlier volumes were flagged and included as spike regressors during the denoising procedure ^71,72^.

Additional denoising on the rsfMRI data was performed using the CONN toolbox (v18a: www.nitrc.org/projects/conn). Temporal and confounding factors were regressed from each voxel BOLD timeseries and the resulting residual timeseries were filtered using a temporal band-pass filter of 0.008-0.09 Hz to examine the frequency band of interest and to exclude higher frequency sources of noise such as heart rate and respiration. For noise reduction, we used the anatomical component-based noise correction method aCompCor, which models the influence of noise as a voxel-specific linear combination of multiple empirically estimated noise sources by deriving principal components from noise regions of interest and including them as nuisance regressors in the first level general linear model (GLM)^73^. Residual head movement parameters (three rotations, three translations, and six parameters representing their first-order temporal derivatives) and signals from WM and CSF, and spike regressors from motion detection were regressed out during the computation of functional connectivity maps.

First-level ROI-to-ROI functional connectivity analysis was performed using the CONN toolbox. For this analysis, we used 12 FreeSurfer ROIs that included unilateral amygdala, fusiform gyrus/perirhinal cortex, and inferior temporal gyrus as part of the AT network and retrosplenial cortex, parahippocampal cortex, and precuneus as part of the PM network (Figure 1). Semi-partial correlations were used for these first-level analyses to determine the unique variance of each (unilateral) seed, controlling for the variance of all other seed regions entered into the same model. For each participant, the rsfMRI time series within each of the ROIs was extracted and the mean time series was computed. Then, the cross-correlation of each ROI’s time course with every other ROI’s time course was computed, creating a 12 × 12 correlation matrix for each subject. Correlation coefficients (i.e., graph edges) were converted to z-values using Fisher’s r-to-z transformation^38^. As in previous studies^11–13^, the diagonal of the matrix was removed and negative correlations were set to zero as we were mainly interested in positive connections^39^. We also performed the same analyses with inclusion of both positive and negative correlations and observed similar results. Finally, AT and PM network segregation were calculated as the difference in mean within-network FC and mean between-network FC divided by the mean within-network FC^13^.

### PET data processing

As part of our standard PET preprocessing procedure, 1.5T structural MRI data were preprocessed with FreeSurfer to derive ROIs in subject native space. These ROIs were then used for the calculation of PiB-PET global distribution volume ratio (DVR) and region-specific, partial volume corrected (PVC ^40^) FTP standardized uptake value ratio (SUVR) measures. FTP images were processed with SPM12. Images were realigned, averaged, and coregistered to each participant’s 1.5T structural MRI scan. SUVR images were calculated by averaging the mean tracer uptake over the 80-100 minute data and normalized by an inferior cerebellar gray reference region ^74^. The mean SUVR of each (FreeSurfer segmented) ROI was extracted from the native space images. This data was then partial volume corrected using a modified Geometric Transfer Matrix approach ^75^ as previously described ^40^. The weighted mean (by region size), partial volume corrected FTP SUVR of all AT (amygdala, FuG/perirhinal cortex, ITG) and PM (RSC, PHC, and precuneus) ROIs were used in subsequent analyses. Tau positivity was defined as the mean SUVR in a BraakIII-IV composite ROI (cut-off 1.26) that included regions from both AT (amygdala, FuG, and ITG) and PM (PHC, RSC) systems. *APOE* was not related to segregation (*ps* > 0.63); therefore, we did not control for *APOE* in these analyses.

PiB images were also processed with SPM12. Images were realigned, averaged across frames from the first 20 minutes of acquisition, and coregistered to each participant’s 1.5T structural MRI image. DVR values for PiB-PET images were calculated with Logan graphical analysis over 35-90 minute data and normalized by a cerebellar gray matter reference region ^76,77^. Global PiB was calculated across cortical FreeSurfer ROIs as previously described^41^, and a threshold of 1.065 was used to classify participants into Aβ− and Aβ+ groups^42^. One participant was missing PiB DVR data, and therefore was excluded from all analyses involving measures of Aβ.

### Statistical analyses

Statistical analyses were conducted using R (http://www.R-project.org/) and SPSS (SPSS Inc., Chicago IL) software. Independent sample t-tests were used to test for age group differences in within- and between-network FC and segregation. Pearson correlations and multiple regression models were used to assess the relationship between segregation, Aβ and tau, and cognitive performance. To assess the relationship between tau and segregation, we first examined the relationship across all OA, regardless of Aβ or tau status. We then split the group into Aβ− and Aβ+ subgroups to determine whether there was an interaction of Aβ pathology and tau on segregation. We used the same procedure to assess the relationship between Aβ and network segregation. Similarly, to assess the relationship between segregation and cognition, we first examined this relationship across all OA, regardless of Aβ or tau status. We then split the group into Aβ− and Aβ+ subgroups to determine whether there was an interaction of Aβ pathology and segregation on cognitive performance. As a follow-up analysis, we also split the groups into tau- and tau+ groups to confirm that the results remained the same. As the results did not change depending on whether we split the groups by Aβ or tau status, we report the results in the present study by splitting the group by Aβ status.

Longitudinal cognitive measures were modelled using linear mixed-effects regression with a random intercept and slope using the lme4 package in R v3.6.3 (www.r-project.org). In order to examine the relationship between baseline segregation, Aβ and tau, and change in cognitive performance, the models included two-way interactions between baseline segregation and time, baseline Aβ and time as well as tau and time. We were specifically interested in the segregation × time interaction to determine whether baseline segregation was associated with longitudinal episodic memory decline. As a control analysis, we also examined change in working memory and executive function performance over time using the same model.

All predictor variables were standardized before entered into the model. All models were adjusted for age and sex, and years education (for models including cognitive measures). All statistical analyses used a two-tailed level of 0.05 for determining statistical significance. Reported *p*-values were not corrected for multiple comparisons.

## Author Contributions

Conceptualization, K.C. and W.J.J.; Methodology & Software: K.E.C., J.N.A., X.C., A.M., T.M.H., S.L., and S.B.; Formal Analysis: K.E.C., J.N.A., X.C., A.M., T.M.H., and S.B.; Writing – Original Draft, K.C. and W.J.J.; Writing – Review & Editing, all authors.

## Supplemental Material

### AT and PM networks are less segregated with older age regardless of analysis approaches

The relationship between age group and segregation was assessed across multiple analysis approaches related to matrix thresholding, bivariate vs. semi-partial correlations, various network metrics of intersystem relationships, and network labeling.

To assess the influence of matrix thresholding on age group differences in network segregation, this supplementary analysis was identical to the original analysis (in the main text) except we retained both positive and negative correlations in each subject’s z-matrix (Figure S1A). Similarly, to assess the influence of type of correlation analysis on age group differences in network segregation, this supplementary analysis was identical to the original one except we calculated bivariate correlations instead of semi-partial correlations (Figure S1B).

To examine the influence of network labeling on age group differences in network segregation, we used two different labeling schemes (in addition to our original one). First, we used the Brainnetome Atlas^1^, using all ROIs from amygdala (4 ROIs), FuG (6 ROIs), ITG (14 ROIs), PHC (12 ROIs), RSC (4 ROIs), and precuneus (8 ROIs). To create these ROIs, we produced 4mm-radius spheres centered around each MNI coordinate from the literature (Figure S1C). To create AT and PM networks based on the same AT and PM regions used in one of the seminal papers defining AT and PM systems^2^, we added 4 FreeSurfer ROIs each to our original AT and PM networks. These additional FreeSurfer regions included bilateral lateral orbitofrontal cortex and temporal pole for the AT network and medial orbitofrontal and posterior cingulate cortices for the PM network, creating a total of 10 ROIs for each network (Figure S1D).

To assess the influence of various network metrics of intersystem relationships, we used the Brain Connectivity Toolbox in Matlab to calculate participation coefficient and modularity values for each subject. The participation coefficient of a given ROI measures to what extent an ROI interacts with ROIs in other networks in relation to the total number of connections it contains in its own network. Each subject’s participation coefficient value was calculated from their respective z-matrix. Participation coefficients for each ROI were computed based on the following formula:

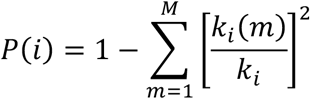

where *k*_*i*_(*m*) is the weighted connections of ROI *i* with nodes in network *m* and *k*_*i*_ is the total weighted connections ROI *i* exhibits. Thus, higher participation coefficient values indicate proportionally greater communication with ROIs in other networks (Figure S1E).

Similar to network segregation, modularity assesses the strength of module (i.e., network) segregation. Specifically, the modularity index (Q) compares the observed intra-module functional connectivity with that which is expected by chance. Thus, higher modularity values reflect stronger separation of the system’s modules. The modularity index is formally defined as:

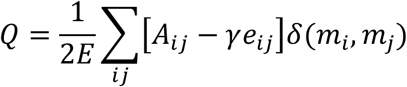

Where *E* is the number of graph connections (i.e., edges), *A* is the adjacency matrix, *γ* is the resolution parameter, *e* is the null model, and *δ* is an indicator that equals 1 if ROIs *i* and *j* belong to the same module and 0 otherwise (Figure S1F).

**Figure S1.**
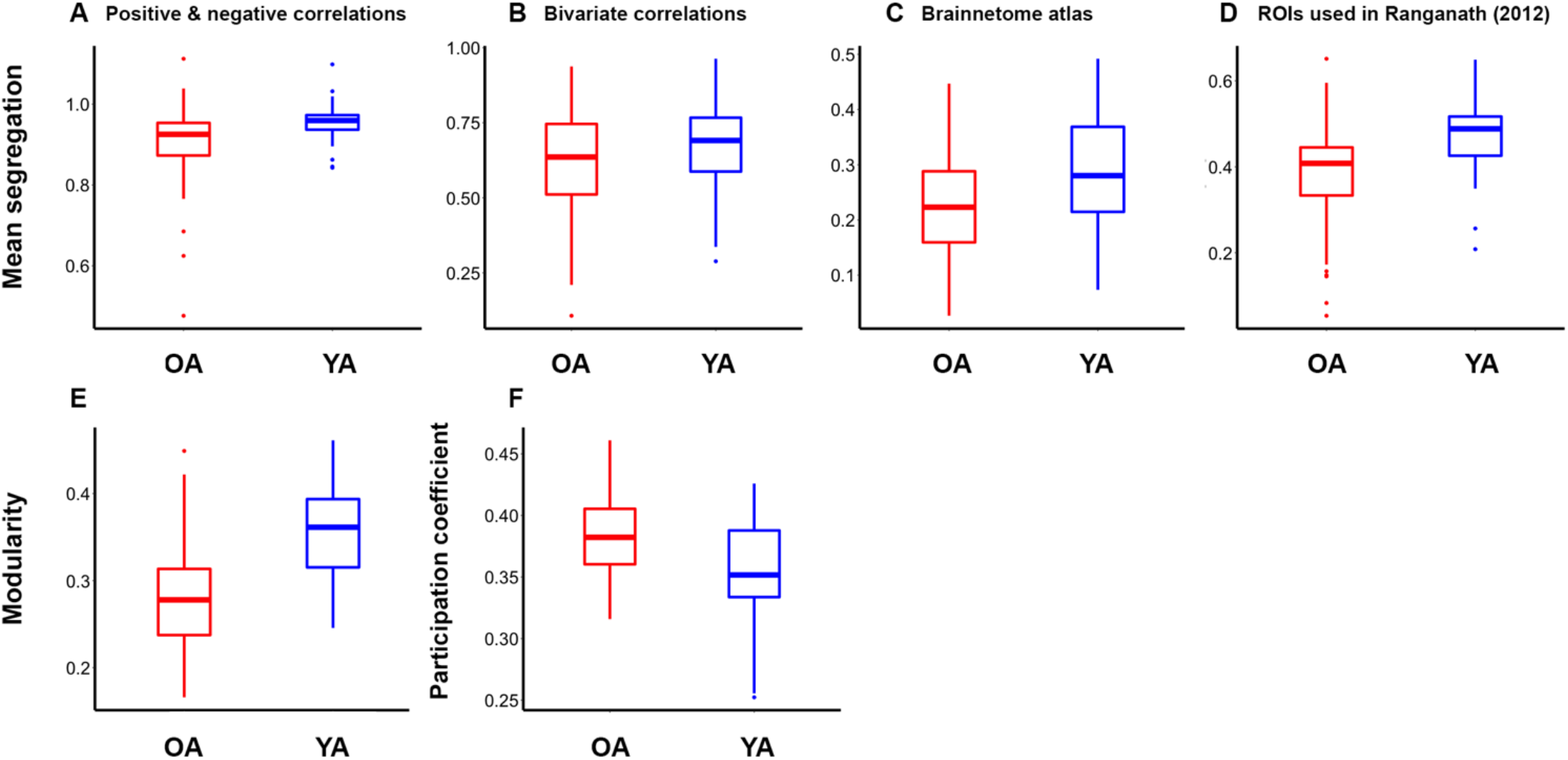
Older adults show significantly less segregated networks compared to younger adults across multiple analysis approaches related to A) matrix thresholding: inclusion of positive and negative correlations (*t* = 2.8, *p* = 0.006), B) bivariate correlations (*t* = 2.4, *p* = 0.019), network labeling: C) Brainnetome Atlas (*t* = 3.4, *p* = 0.001) and D) inclusion of 20 FreeSurfer ROIs used in Ranganath et al. 2012. (*t* = 5.7, *p* < 0.001), and various network metrics of intersystem relationships: E) participation coefficient (*t* = 4.7, *p* < 0.001), and F) modularity (*t* = 7.9, *p* < 0.001).

### AD pathology moderates the association between segregation and episodic memory in AT and PM networks

To examine the association between network segregation and episodic memory in the main text, we computed a single segregation measure by averaging the AT and PM segregation values because our episodic memory composite measure included both object- and spatial-related memory domains. Figure S2 below shows that the results were essentially the same in both the AT and PM networks separately.

**Figure S2.**
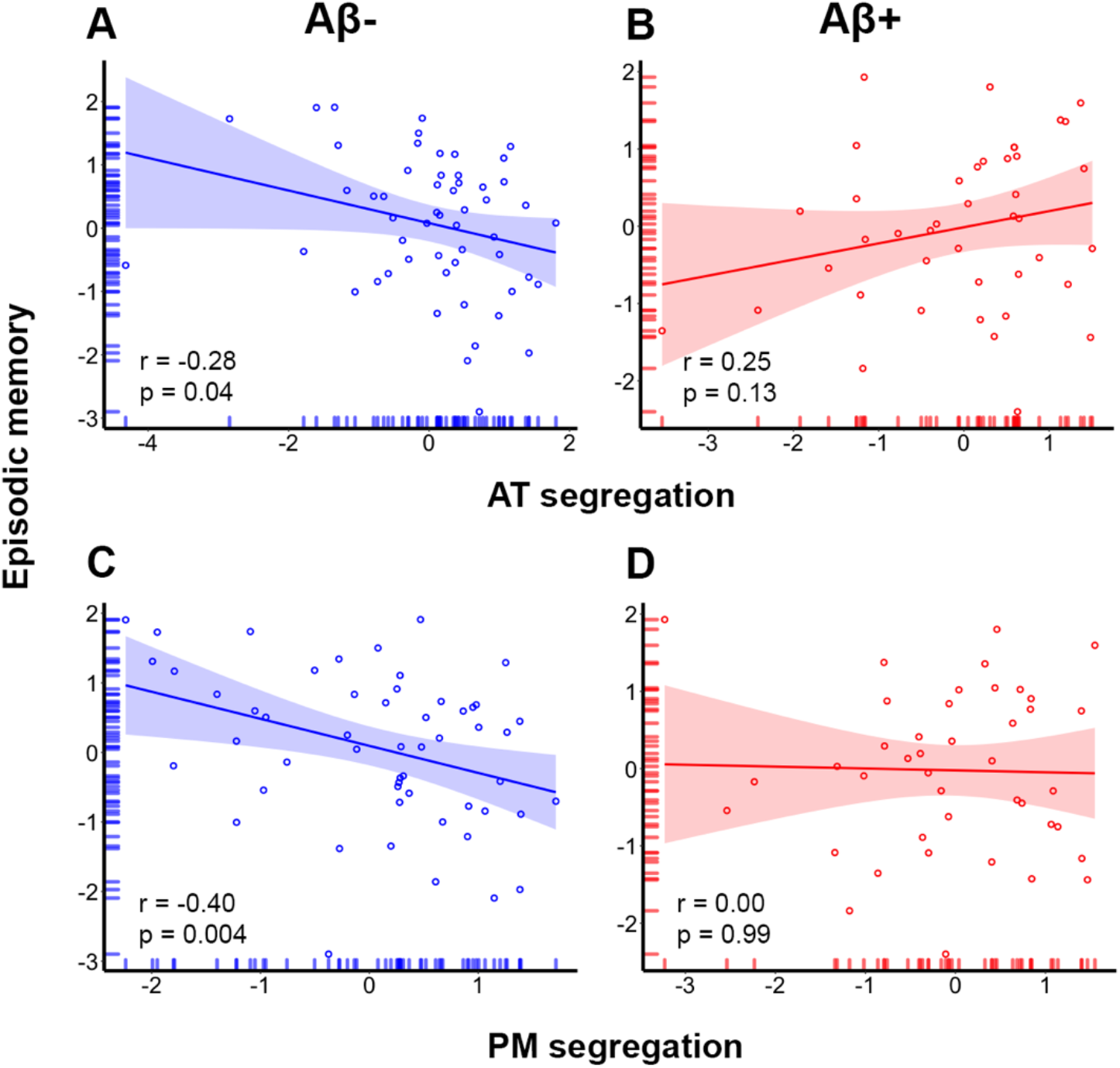
Alzheimer’s disease pathology moderates the association between network segregation and episodic memory performance in AT and PM systems. (**A**) Less segregated AT networks are associated with better performance in Aβ− older adults whereas **(B)** AT segregation is not associated with performance in Aβ+ older adults. (**C**) Similarly, less segregated PM networks are associated with better performance in Aβ− older adults whereas **(D)** PM segregation is not associated with performance in Aβ+ older adults.

### Baseline segregation predicts longitudinal memory decline using various measures of pathology

In order to examine the relationship between baseline segregation, baseline Aβ and tau, and change in cognitive performance, we used a linear mixed model that included two-way interactions between baseline segregation and time, global Aβ and time, and tau and time. In the main text, we report the results using continuous measures of global Aβ and Braak_III-IV_ tau as they retain more statistical power in the model. The results were very similar whether we used Braak_III-IV_ tau (Table 3 in main text), AT-tau (Table S1) or PM tau (Table S2) and whether we used dichotomous (Table S3) or continuous Aβ and tau (Table 3 in main text) in the model. All models were adjusted for age, sex, and education.

**Table S1.** Linear mixed model results for segregation and pathology (including AT-tau) predicting longitudinal episodic memory change.

| Predictor | Estimate | p |
| --- | --- | --- |
| Age | -0.20 | 0.02 |
| Sex | -0.003 | 0.97 |
| Education | 0.13 | 0.12 |
| Segregation | -0.23 | 0.007 |
| Time | -0.07 | 0.051 |
| AT-tau | -0.11 | 0.4 |
| Aβ | 0.03 | 0.79 |
| AT-tau x Time | -0.19 | <0.001 |
| Aβ x Time | 0.007 | 0.87 |
| Segregation x Time | 0.07 | 0.049 |

**Table S2.** Linear mixed model results for segregation and pathology (including PM-tau) predicting longitudinal episodic memory change.

| **Predictor** | **Estimate** | **p** |
| --- | --- | --- |
| Age | -0.20 | 0.03 |
| Sex | 0.02 | 0.84 |
| Education | 0.13 | 0.13 |
| Segregation | -0.18 | 0.036 |
| Time | -0.07 | 0.071 |
| PM-tau | -0.11 | 0.29 |
| Aβ | 0.03 | 0.79 |
| PM-tau x Time | -0.08 | 0.073 |
| Aβ x Time | -0.05 | 0.25 |
| Segregation x Time | 0.07 | 0.05 |

**Table S3.** Linear mixed model results for segregation and (dichotomous) pathology predicting longitudinal episodic memory change.

| **Predictor** | **Estimate** | **p** |
| --- | --- | --- |
| Age | -0.20 | 0.02 |
| Sex | 0.007 | 0.94 |
| Education | 0.13 | 0.15 |
| Segregation | -0.18 | 0.032 |
| Time | -0.06 | 0.09 |
| Tau-status | 0.009 | 0.93 |
| Aβ-status | 0.03 | 0.78 |
| Tau-status x Time | -0.09 | 0.03 |
| Aβ-status x Time | -0.01 | 0.78 |
| Segregation x Time | 0.07 | 0.05 |

